# Imaging the master regulator of the antioxidant response in non-small cell lung cancer with positron emission tomography

**DOI:** 10.1101/2023.12.16.572007

**Authors:** Hannah E. Greenwood, Richard S. Edwards, Will E. Tyrrell, Abigail R. Barber, Friedrich Baark, Muhammet Tanc, Eman Khalil, Aimee Falzone, Nathan P. Ward, Janine M. DeBlasi, Laura Torrente, David R. Pearce, George Firth, Lydia M. Smith, Oskar Vilhelmsson Timmermand, Ariana Huebner, Madeleine E. George, Charles Swanton, Robert E. Hynds, Gina M. DeNicola, Timothy H. Witney

## Abstract

Mutations in the NRF2-KEAP1 pathway are common in non-small cell lung cancer (NSCLC) and confer broad-spectrum therapeutic resistance, leading to poor outcomes. The cystine/glutamate antiporter, system x_c_^−^, is one of the >200 cytoprotective proteins controlled by NRF2, which can be non-invasively imaged by (*S*)-4-(3-^18^F-fluoropropyl)-ʟ-glutamate ([^18^F]FSPG) positron emission tomography (PET). Through genetic and pharmacologic manipulation, we show that [^18^F]FSPG provides a sensitive and specific marker of NRF2 activation in advanced preclinical models of NSCLC. We validate imaging readouts with metabolomic measurements of system x_c_^−^ activity and their coupling to intracellular glutathione concentration. A redox gene signature was measured in patients from the TRACERx 421 cohort, suggesting an opportunity for patient stratification prior to imaging. Furthermore, we reveal that system x_c_^−^ is a metabolic vulnerability that can be therapeutically targeted for sustained tumour growth suppression in aggressive NSCLC. Our results establish [^18^F]FSPG as predictive marker of therapy resistance in NSCLC and provide the basis for the clinical evaluation of both imaging and therapeutic agents that target this important antioxidant pathway.

## Introduction

To provide defence against oxidative stress and maintain redox homeostasis, cells upregulate a myriad of different detoxifying enzymes and transporters. This antioxidant program is mediated in part by the transcription factor nuclear factor-erythroid 2 p45-related factor two (NRF2, *NFE2L2*). Under normal conditions, NRF2 protein levels are controlled by its negative regulator, Kelch-like ECH-associated protein 1 (KEAP1). KEAP1 binds to NRF2 in the cytosol, recruiting the Cullin-3 (Cul3) E3 ubiquitin ligase complex which ubiquitinates NRF2, leading to its proteasomal degradation. Under conditions of oxidative stress, elevated levels of reactive oxygen species (ROS) oxidise key cysteine residues in KEAP1 (C226, C273 and C288), altering its conformation, thereby impairing the degradation of NRF2. Newly translated NRF2 subsequently accumulates in the cell.^1^

Cancer cells exploit NRF2 activation as a pro-survival technique to overcome insult from ROS^2, 3^. NRF2 stabilisation is facilitated by either gain of function mutations in NRF2, or loss of function mutations in KEAP1. Additionally, epigenetic alterations, including CpG *KEAP1* promoter hypermethylation, mutations in the Cul3 ligase complex, and oncogene-dependent transcriptional upregulation, alter KEAP1 function^3, 4^. Following its stabilisation, NRF2 directly upregulates enzymes that control glutathione (GSH) biosynthesis; enzymes involved in phase 2 drug metabolism, such as NAD(P)H quinone oxidoreductase 1 (NQO1) and glutathione S-transferase (GST)^5^; drug efflux pumps, including ATP-binding cassette (ABC) transporters and multidrug resistance (MDR) proteins^6^; homologous DNA repair proteins^7^; and anti-apoptotic proteins, such as Bcl-2^8^. Unsurprisingly, hyperactivation of NRF2 results in resistance to a broad range of traditional and emerging therapies, including chemotherapy^9^, radiotherapy^10^, immunotherapy^11^, and more recently, escape from KRAS G12C inhibitor treatment^12^.

Activating mutations in NRF2/KEAP1 are found in approximately a third of non-small cell lung cancer (NSCLC) patients^13, 14^, with NRF2 activation associated with a worse prognosis in patients treated with first-line therapy^3, 15^. Ten-year survival rates for NSCLC are stuck at approximately 5%^16^, meaning there is an urgent clinical need to identify patients with poor prognosis and develop therapeutics that exploit specific vulnerabilities in these aggressive tumour subtypes. One of the >200 cytoprotective proteins directly regulated by NRF2 is system x_c_^−17^. System x_c_^−^ is a heterodimeric transporter that consists of two subunits: the functional transporter and light chain xCT (*SLC7A11*), which confirms substrate specificity, and CD98hc (*SLC3A2*), the membrane protein common to many amino acid transporters which is responsible for membrane localisation and activity^18, 19^. System x_c_^−^ functions to exchange the intracellular amino acid glutamate for extracellular cystine, which, following reduction to cysteine, provides the rate-limiting precursor for *de novo* biosynthesis of GSH^20^, the body’s most abundant antioxidant.

We have previously evaluated the positron emission tomography (PET) radiotracer (*S*)-4-(3- ^18^F-fluoropropyl)-ʟ-glutamate ([^18^F]FSPG) as a specific measure of system x_c_^−^ activity in living subjects^21–23^. In the current study, we investigated whether tumour-intrinsic KEAP1/NRF2 mutations could be non-invasively imaged by [^18^F]FSPG PET imaging and whether aberrant system x_c_^−^ expression could be exploited for treatment of refractive NSCLC tumours. Across a panel of NSCLC lines and through pharmacological and genetic manipulation of NRF2, we found that [^18^F]FSPG provided a sensitive readout of NRF2 activation. To assess the translational relevance of our findings, we examined [^18^F]FSPG retention in mice bearing orthotopically grown NRF2-high and NRF2-low NSCLC tumours, in the Kras^G12D/+^; p53^flox/flox^ adenocarcinoma model with and without Nrf2^D29H^ mutations, and in patient-derived human xenografts. [^18^F]FSPG retention was significantly higher in tumours with constitutively active NRF2 activity across these varied animal models of NSCLC. Having used imaging to identify NRF2-high NSCLC tumours, we finally showed that a novel antibody-drug conjugate (ADC) targeting system x_c_^−^ induced prolonged tumour growth suppression in drug-resistant cancer. Taken together, our study reveals that NRF2 activation provides a metabolic susceptibility for targeted imaging and treatment.

## Methods

### Cell lines and cell culture

Human NSCLC lines A549, H460, H1944, H1792, H23, H1299, H1975 and H1650 were purchased from LGC Ltd. H460 FLuc were purchased from PerkinElmer. H1299 FLuc were purchased from AMS biotec. A549 NRF2 knockout (KO), A549 KO-restored (KO-R), H1299 T80K and H1299 PLX317 were previously described ^24, 25^. To pharmacologically activate NRF2 expression, cells were treated with 100 nM of the KEAP1 inhibitor, KI696, for 24 h. All cells were cultured in RPMI media (Sigma-Aldrich), supplemented with 10% foetal bovine serum (ThermoFisher Scientific) and 100 U.mL^−1^ penicillin and 100 µg.mL^−1^ streptomycin (Sigma-Aldrich), maintained at 37 °C and 5% CO_2_. Cell lines were tested monthly for mycoplasma infection (Eurofins).

Cells were seeded for 24 h prior to experimental endpoint in 6-well plates in 2 mL of media. All cells were seeded at 1.5 × 10^5^/mL, except for H1944 and H1792, which were seeded at 2.75 × 10^5^/mL; H1650, which was seeded at 1.75 × 10^5^/mL; and H23 and H1975, which were seeded at 2.5 × 10^5^/mL. For KI696 treatments, H1650, H23, H1975 and H1299 were seeded at 1.0 × 10^5^/mL, 1.0 × 10^5^/mL, 1.25 × 10^5^/mL and 0.75 × 10^5^/mL, respectively 2 mL of media 24 h prior to treatment.

### Western blotting

Western blot analysis was carried out using an iBind Flex system (ThermoFisher Scientific), as described previously ^26^. For cell lysate collection, cells were seeded for 24 h prior to harvesting, at densities described above. Briefly, cells were placed on ice, washed three times with ice-cold PBS and lysed in 100 µL RIPA buffer containing 1× protease and phosphatase inhibitors (ThermoFisher Scientific). Collected lysates were then centrifuged at 15,000 × *g* at 4 °C for 10 min. Cell debris was removed and the supernatant aliquoted to avoid freeze-thaw cycles.

For *ex vivo* sample preparation, tissues were dissected immediately following sacrifice, snap frozen in liquid nitrogen, and stored at -80 °C. Frozen tissues were added to pre-chilled Lysing Matrix tubes containing 1.4 mm ceramic beads (MP Biomedicals) and 1 mL RIPA buffer containing 1 × protease and phosphatase inhibitors. Samples were lysed by rapid shaking using a high-speed benchtop reciprocating homogenizer, cooled to 4 °C (Precellys 24 homogenizer, Bertin Instruments). The lysates were centrifuged at 15,000 × *g* at 4 °C for 10 min and the supernatant collected for analysis. The protein content in each lysate was determined using a BCA assay kit (ThermoFisher Scientific) and 20 µg of protein was loaded into each well.

Blots were probed for xCT, NRF2 and NQO1 (1:1000 dilution; Cell Signaling Technology). Actin was used as a loading control for all experiments, with a horseradish peroxidase (HRP) linked anti-rabbit IgG secondary antibody (1:200 dilution; Cell Signaling Technology). After antibody incubations, membranes were removed from the iBind Flex system and washed in 50 mL of tris-buffered saline with 0.1% Tween 20 (TBST), five times for 5 min on a shaker (Stuart gyratory rocker SSL3). To visualize proteins, 4 mL of Amersham™ ECL Prime Western Blotting Detection Reagent (GE HealthCare) was added to each membrane in the dark for 1 min. Images were taken using an iBright CCD camera (Invitrogen). Images were always acquired within the linear range of the camera to prevent the overexposure of any blots.

### ^18^F-FSPG radiosynthesis

^18^F-FSPG was synthesised using a GE Fastlab according to previously-published methodology^27^.

### Radiotracer uptake and efflux

Cells were seeded into 6-well plates 24 h prior to uptake and efflux studies (see previous). For all cell uptake and efflux experiments, 0.185 MBq/mL of [^18^F]FSPG was added to wells (total volume 1 mL/well) and incubated for 1 h. Cells were maintained at 37 °C and 5% CO_2_ throughout the uptake or efflux experiment. For uptake studies, plates were removed from the incubator after a 60 min incubation and placed on ice. Cells were washed three times with ice-cold PBS before addition of 0.5 mL of RIPA buffer. 0.3 mL of cell lysate was taken for counting and the remaining 0.2 mL was used for protein quantification (BCA). For efflux experiments, following 60 min uptake, exogenous radioactivity was removed, with cells washed three times in room temperature PBS, and fresh media was added for 20, 40 or 60 min before being placed on ice and processed as above. The amount of radioactivity in the samples was determined in a gamma counter (1282 Compugamma, LKB Wallac) and expressed as a percentage of the administered dose per mg protein.

### ROS quantification

ROS were detected in human NSCLC lines using the cell permeable fluorophore CellROX Green (ThermoFisher Scientific). Cells were seeded in 6-well plates densities stated above in 2 mL media 24 h prior to performing the assay. A final concentration of 1 μmol/L CellROX Green reagent was added to each well and incubated for 30 min at 37 °C, protected from light. Cells were then washed with PBS, harvested using 0.05% trypsin-EDTA and suspended in 1 mL of ice-cold Hanks balanced salt solution. The cell suspension was passed through a 35 μm filter and kept on ice prior to analysis on BD FACS Melody flow cytometer (488 nm laser and 527/32 bandpass filter; BD Biosciences). 20,000 single cell events were recorded per sample with the data gated post-acquisition based on forward (FS) and side scatter (SS) profiles to include only single cell events and to exclude cellular debris.

### Glutamate quantification

Intracellular glutamate concentrations were determined using a glutamate colorimetric assay kit, following the manufacturers guidelines (Biovision). Briefly, cells seeded in a 6-well plate were trypsinised, washed three times with ice-cold PBS and resuspended in 200 µL of assay buffer before sonication on ice (Soniprep 150, MSE). Lysates were then centrifuged at 15,000 × *g* for 10 min at 4 °C and the supernatant taken for analysis. Total intracellular glutamate was normalized to protein concentration.

### GSH

Cells were seeded into 6-well plates as described above. Total GSH was determined using luminescence-based quantification (GSH/GSSG-Glo Assay, Promega). Cells were washed in ice-cold PBS, lysed in assay buffer, and centrifuged at 15,000 × *g* at 4°C for 10 min. For *in vivo* studies, tumour tissue was dissected, snap frozen in liquid nitrogen, and stored at −80°C. When required, tissue was placed into lysis Matrix tubes containing 1.4-mm ceramic beads (MP Biomedicals) and assay buffer (Promega). Tumour samples were lysed by rapid shaking using a high-speed benchtop reciprocating homogenizer (Fastprep-24 Sample Preparation Instrument; MP Biomedicals). The lysates were centrifuged at 15,000 × *g* at 4°C for 10 min and the supernatant collected for analysis. For all studies, 5 μL of supernatant along with 5 μL of GSH standards (1–100 μmol/L) were added to white 96-well plates and total GSH determined according to manufacturer’s instructions. Total intracellular GSH was normalized to protein concentration.

### Cystine

5 µL of conditioned media from cultured NSCLC cells subject to the indicated treatment was extracted in 195 µL of ice-cold 82% methanol supplemented with 25 mM N-ethylmaleimide, 10 mM ammonium formate, and 10 µM [D_4_]cystine (Cambridge Isotope Laboratories, DLM-1000-1). Following a 30 min incubation at 4 °C, extracts were cleared by centrifugation at 17,000 × *g* for 20 minutes at 4 °C. 5 µL of cleared extract was then analysed by LC-MS according to an established protocol^24^. Briefly, a Vanquish UPLC system was coupled to a Q Exactive HF (QE-HF) mass spectrometer equipped with HESI (Thermo Fisher Scientific) for chromatographic metabolite separation. Samples were then run on an Atlantis Premier BEH Z-HILIC VanGuard FIT column, 2.5 µm, 2.1 mm × 150mm (Waters). The mobile phase A was 10 mM (NH_4_)_2_CO_3_ and 0.05% NH_4_OH in H_2_O, while mobile phase B was 100% ACN. The column chamber temperature was set to 30°C. The mobile phase condition was set according to the following gradient: 0-13 min, 80% to 20% of mobile phase B; 13-15min, 20% of mobile phase B. The ESI ionization mode was negative, and the MS scan range (m/z) was set to 65-975. The mass resolution was 120,000 and the AGC target was 3 × 10^6^. The capillary voltage and capillary temperature were set to 3.5 kV and 320°C, respectively. The cystine and [D_4_]cystine peaks were manually identified and integrated with EL-Maven (Version 0.11.0) by matching to an in-house library. Cystine concentrations were calculated based on the 10 µM [D_4_]cystine internal standard peak for each sample. Cystine consumption rates were calculated based on the change in media cystine levels following the 4 h culture duration and normalized to cell density.

### Animal studies

All animal experiments performed in the U.K. were in accordance with the United Kingdom Home Office Animal (scientific procedures) Act 1986 and received local approval. Experiments were performed unblinded.

### Orthotopic tumour cell implantation

1 × 10^6^ H460 FLuc cells or 5 × 10^6^ H1299 FLuc were administered by a non-invasive intratracheal technique into the lungs of female NSG mice aged 6-9 weeks (Charles River Laboratories) described previously ^21^. Briefly, mice were anaesthetized with isoflurane (2-2.5% in O_2_) and transferred onto a vertical board where they were suspended in an upright position by their upper incisors. Using a Leica M125 stereomicroscope (Leica Microsystems), the tongue was moved to one side to expose the vocal cords and the entrance to the trachea, where a plastic 20G i.v. catheter, attached to a 1 mL syringe was inserted for cell administration, suspended in 50 µL of PBS. Tumour growth was monitored through bioluminescence imaging.

### Bioluminescence imaging

Orthotopic H460 FLuc and H1299 FLuc tumours were monitored through bioluminescence imaging using an IVIS Spectrum *in vivo* imaging system (PerkinElmer). Images were acquired 4 h post cell inoculation to confirm successful delivery of cells to the lung. Mice were subsequently imaged once a week for the first three weeks and then every 3-4 days thereafter. Mice were anesthetized with isoflurane (1-2 % in O_2_) and injected i.p. with 150 mg/kg firefly luciferin (Promega) before being transferred to the IVIS Spectrum camera and maintained at 37 °C. Images were acquired until the luminescent signal plateaued ∼20 min p.i. of luciferin, ensuring maximum tumour signal was reached (exposure time 1-60 s, binning 2-8, FOV 23 cm, f/stop 1, no filter). Tumour growth was monitored until experimental endpoint. For signal quantification, images were analysed using Living Image software (PerkinElmer). A region of interest was drawn around the entire thorax and total photon flux was measured (photon/sec). Once the bioluminescent signal reached ∼5 × 10^7^ photons/s/cm^3^, mice were selected for PET/CT imaging.

### Genetically engineered mice

LSL-Kras^G12D/+^; p53^flox/flox^; Nrf2^D29H/+^ mice were described previously^28^. Mice were housed and bred in accordance with the ethical regulations and approval of the University of South Florida Institutional Animal Care and Use Committee (protocol numbers: IS00005814M and IS00003893R). All mice were maintained on a mixed C57BL/6 genetic background. Lung tumours were induced by intranasal installation of 2.25 × 10^7^ PFU adenoviral-Cre (University of Iowa) as previously described^28^. Adenoviral infections were performed under isofluorane anesthesia. Following infection, mice were shipped to the U.K. for imaging studies.

### Analysis of transcriptomic data

Bulk RNA sequencing data were obtained from the TRACERx 421 study^29^. Once normal tissue and lymph node samples were removed, this comprised 882 samples from 344 NSCLC patients (median = 2 samples/patient; interquartile range: 2-3) with 279 having multi-region sampling. Z-score was calculated using the equation z = (x-μ)/σ, where x is the raw score, μ is the population mean, and σ is the population standard deviation.

### Patient-derived xenografts

Ethical approval to generate patient-derived models was obtained through the TRACERx clinical study (REC reference: 13/LO/1546; NCT01888601). PDX models were developed by subcutaneous injection of minced primary, surgically resected NSCLC tumour tissue in NOD/SCID/IL2Rg^-/-^ (NSG) mice, as previously described^30^. PDX models were passaged twice prior to cryopreservation and shipping to King’s College London for imaging experiments. Driver mutation profiles of PDX models were derived from whole-exome sequencing of both primary tumours and PDX models, and are reported according to the definition previously described^30^. Cryopreserved tumour samples were thawed in a 37 °C water bath, transferred to a sterile 10 cm dish, and minced in fresh RPMI medium containing 1× penicillin/streptomycin. On ice, tumour pieces were then added to an Eppendorf containing 180 μL of Matrigel (Corning) before being injecting into the upper flank of anesthetised female NSG mice (6-8 weeks) using a 19G needle. Mice were monitored daily and selected for [^18^F]FSPG PET/CT imaging when tumours reached ∼150 mm^3^.

### PET/CT imaging

Mice received a single bolus i.v. injection through a tail vein cannula of ∼3 MBq [^18^F]FSPG in 100 µL PBS. 40 min post-injection, static PET imaging scans were acquired for 20 min on a Mediso NanoScan PET/CT system (1-5 coincidence mode; 3D reconstruction; CT attenuation-corrected; scatter corrected). CT images were acquired for anatomical visualization and attenuation correction (720 projections; semi-circular acquisition; 55 kVp; 600 ms exposure time). Animals were maintained under anaesthesia (1-2% isoflurane in O_2_) at 37 °C during tail vein cannulation, radiotracer administration, and throughout the scan. Static image reconstruction was performed using the 3D Tera-Tomo algorithm, with 4 iterations, 6 subsets, a 0.4 mm isotropic voxel size and a binning window of 400-600 keV. VivoQuant software (v 2.5, Invicro Ltd) was used for image quantification of the reconstructed scans. Individual tumour volumes of interest in the lungs of mice were constructed by manually drawing sequential 2D regions of interest (ROI) on the CT images. The radioactivity in each individual lesion was expressed as a percentage of the injected dose per gram of tissue volume (%ID/g).

The percentage of tumour tissue in the lungs of individual mice was determined using the total CT voxel volume. Individual ROIs were manually drawn using VivoQuant from CT images, and the tumour volumes were summed for individual animals. Additionally, a single ROI was drawn around the entire lung to determine total lung volume. Together, the tumour volumes and total lung volume were used to calculate the percentage of tumour tissue present.

### *Ex vivo* tumour autoradiography

Following [^18^F]FSPG PET/CT imaging, tumours were perfused with PBS prior to being embedded and frozen in a 1:1 OCT-PBS mixture using an isopentane bath over liquid nitrogen. The cryopreserved tumours were then sectioned (20 μm) using a Bright 5040 cryotome, with tissue sections thaw-mounted onto Superfrost PLUS glass microscope slides (Menzel-Glaser, Thermo). The slides were then exposed to a storage phosphor screen (Cytiva) and stored in a standard x-ray cassette overnight. The phosphor screen was then imaged using an Amersham Typhoon scanner (Cytiva) and for anatomical reference, hematoxylin and eosin (H&E) staining was performed.

### Immunohistochemistry

Consecutive sections of formalin-fixed, paraffin-embedded tumour-containing lungs were used for immunohistochemical (IHC) analysis of xCT using a VECTOR DAB substrate kit (Vector laboratories) following the manufacturer’s instructions. Slides were deparaffinized in xylene and rehydrated through graded ethanol to water before staining. For antigen unmasking, all sections were treated with 10 mM citrate solution (pH 6.0), and with 3% H_2_O_2_ for the inactivation of endogenous peroxidase. Sections were blocked in 2.5% normal blocking serum from the VECTASTAIN Elite ABC-HRP kit (Vector laboratories), prior to incubation with the avidin and biotin solutions, respectively (Vector laboratories). Sections were incubated with the anti-xCT antibody (1:200 dilution; AgilVax) overnight at 4°C. After washing in TBST, the sections were stained with a biotinylated secondary antibody for 30 min at room temperature. After washing with TBST, sections were incubated with DAB substrate solution, and counterstained with haematoxylin. Images were acquired using a NanoZoomer (Hamamatsu), with representative images shown. Four areas were randomly selected from the tissue sections and the total number of cells and number of stained cells were counted. This was repeated for three individual tumours and the percentage of stained cells was calculated. For the genetically engineered mouse model studies, slides were scanned with the Aperio imager at 20× and the H-score of at least five representative regions/mouse was analyzed with QuPath software^31^. Representative images were captured using the Axio Lab A1 microscope at 40× (Carl Zeiss Microimaging Inc.).

### HM30-tesirine growth inhibition in cells

H460 and H1299 cells were seeded into a 96-well plate at 2,000 cells/well. xCT-targeting ADC was linked to Tesirine and provided by AgilVax (patent: WO2020/227640A1). 24 h post-seeding, cells were treated with increasing concentrations of HM30-Tesirine ranging from 0.02 to 400 nM. After 72 h, media was removed and 100 µL of 0.5 mg/mL MTT solution was added for 4 h. Next, the MTT solution was removed, and formazan crystals were dissolved in 150 µL DMSO before absorbance was measured at 570 nm using a Varioskan Lux (ThermoFisher Scientific).

### HM30-tesirine treatment of H460 tumour-bearing mice

3 × 10^6^ H460 cancer cells in 100 µL PBS were injected subcutaneously into female Balb/C nu/nu mice aged 6-9 weeks (Charles River Laboratories). Tumour growth was monitored using an electronic caliper and the volume calculated using the following equation: volume = ((π/6) × h × w × l), where h, w and l represent, height, width, and length, respectively. When tumours reached ∼70 mm^3^ mice were randomized into HM30-tesirine, cisplatin, and vehicle treatment groups. HM30-tesirine treated animals received three doses of 1.5 mg/kg on days 0, 7 and 14 via an i.p. injection (200 µL). The cisplatin-treated cohort received two doses of cisplatin (5 mg/kg; 200 µL i.p) on days 0 and 4. Vehicle control animals were treated on days 0 and 7. Mice were weighed and tumour volumes measured until humane endpoints were reached.

### Statistics

Statistical analysis was performed using GraphPad Prism (v.8.0). All *in vitro* data was acquired from three or more biological replicates, acquired on separate days. Data were expressed as the mean ± one standard deviation (SD). Statistical significance was determined using unpaired two-tailed Student’s t-test. For analysis across multiple samples, 1-way analysis of variance (ANOVA) followed by t-tests multiple comparison correction (Tukey method) were performed. Kaplan–Meier survival curve statistics were analysed with a log-rank (Mantel–Cox) test. To control the family-wise error rate in multiple comparisons, crude P-values were adjusted by the Holm–Bonferroni method. Differences with *p* values < 0.05 were considered statistically significant in all analyses.

## Results

### NRF2 activation in NSCLC cells increases system x_c_^−^ activity and [^18^F]FSPG cell retention

*SLC7A11* is a NRF2-regulated gene which encodes the glutamate/cystine antiporter xCT, a key component of system x_c_^−^. Here, we examined the role of NRF2 activation in relation to system x_c_^−^ activity, downstream antioxidant capacity, and retention of the system x_c_^−^ substrate [^18^F]FSPG in NSCLC cells grown in culture (**Fig. 1a**). Cell lines harbouring functional *KEAP1* mutations had elevated NRF2 protein expression and increased expression of downstream NRF2 targets compared to cells with wild-type (WT) or silent *KEAP1* mutations (H23; **Fig. 1b**). In these NRF2-high cells, xCT was elevated compared to NRF2-low cells, which was accompanied by a 7.6-fold increase in cystine consumption (8008 ± 2827 nmol/h/100,000 cells vs. 1051 ± 3208 nmol/h/100,000 cells, respectively; *n* = 4; *p* = 0.01; **Fig. 1c**). Conversely, intracellular glutamate concentrations were halved in NRF2-high cells compared to NRF2-low cells (2.0 ± 0.42 nmol/mg protein vs 3.9 ± 1.1 nmol/mg protein, respectively; *n* = 4; *p* = 0.02; **Fig. 1d**), providing evidence that not only xCT expression but transporter activity were increased in NRF2-high NSCLC cells. Both glutamate and cystine are substrates for GSH biosynthesis. In cells with functional *KEAP1* mutations, GSH was increased three-fold compared to NRF2-low cells (1.67 ± 0.46 nmol/mg protein vs. 0.57 ± 0.16 nmol/mg protein, respectively; *n* = 4; *p* = 0.004; **Fig. 1e**). Furthermore, ROS was more than halved in NRF2-high cells compared to NRF2-low cells (*n* = 4; *p* = 0.0003), indicating increased antioxidant capacity in cells that lack functional KEAP1 (**Fig. 1f,g**).

**Figure 1.**
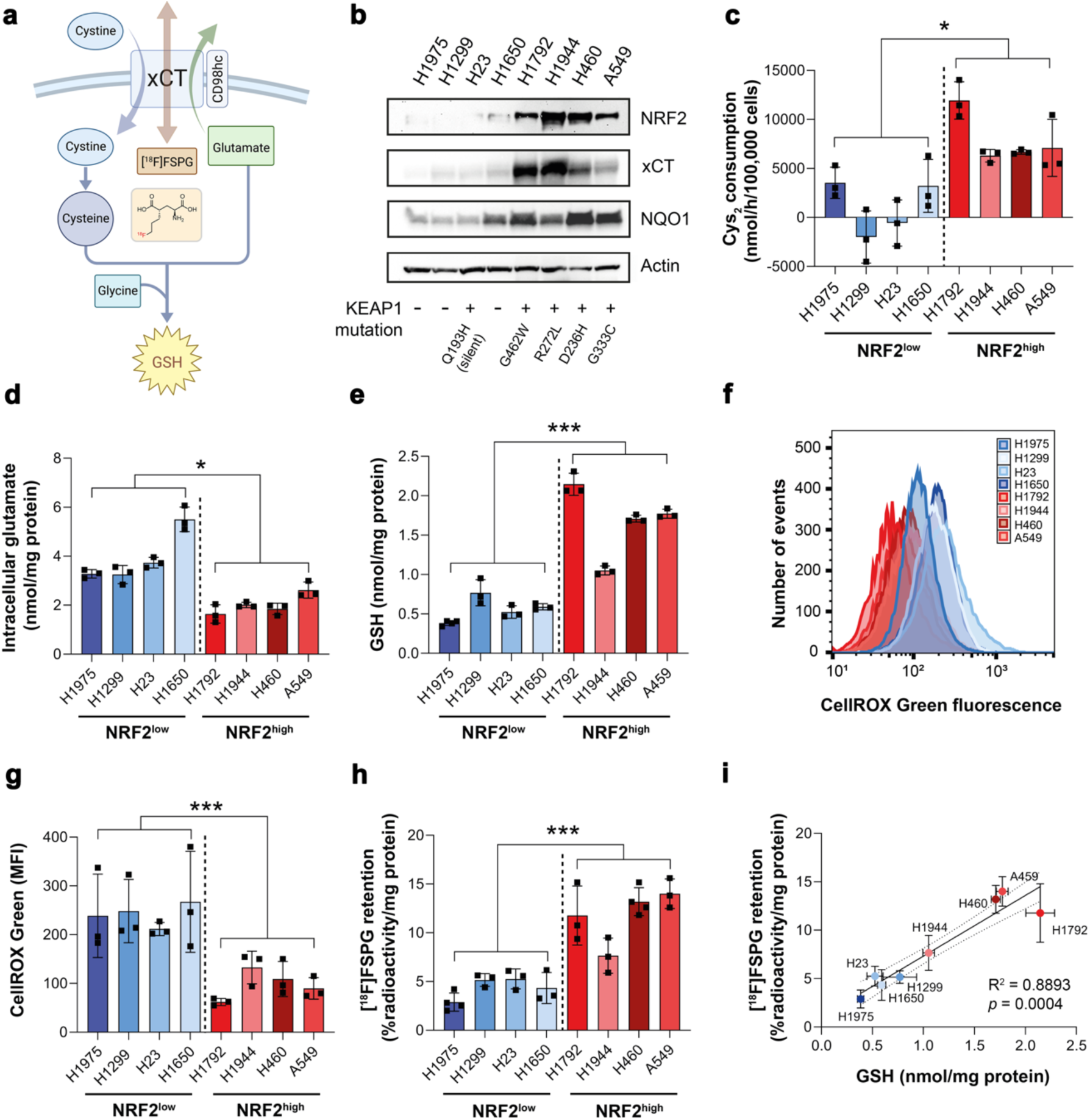
Elevated NRF2 increases xCT expression, system x_c_^−^ activity, and downstream antioxidant capacity, detectable by [^18^F]FSPG. a, Schematic of system x_c_^−^ with its natural substrates cystine and glutamate, and the radiotracer [^18^F]FSPG (structure shown in insert). b, Protein expression of NRF2, xCT and NQO1 in a panel of NSCLC lines and corresponding KEAP1 mutations. Actin was used as a loading control. c, Cystine consumption in NSCLC lines following media replenishment. Cys_2_, cystine. Intracellular glutamate (d) and GSH (e) in NSCLC lines. Flow cytometric measurement of total ROS levels using CellROX Green with representative histograms (f) and median fluorescent intensity (MFI; g) shown. h, Intracellular retention of [^18^F]FSPG. i, Correlation between intracellular GSH and intracellular [^18^F]FSPG accumulation. Broken lines represent the 95% confidence interval of the best fit line. Data are presented as mean ± SD. *, p < 0.05; ***, p < 0.001.

To investigate whether [^18^F]FSPG provides a functional readout of NRF2-status, [^18^F]FSPG retention was evaluated in a panel of eight NSCLC lines. Following 60 min incubation, [^18^F]FSPG retention was 2.6-fold higher in NRF2-high lines compared to NRF2-low lines (11.7 ± 2.8% radioactivity/mg protein vs. 4.4 ± 1.1% radioactivity/mg protein, respectively; *n* = 4; *p* = 0.003; **Fig. 1h**). [^18^F]FSPG cell retention is a product of both radiotracer uptake and efflux, as [^18^F]FSPG can pass bidirectionally across xCT. Although net retention of [^18^F]FSPG was higher in NRF2-high cells, [^18^F]FSPG efflux was also increased in these cells (**Extended Data Fig. 1**), indicating increased system x_c_^−^ activity compared to NRF2-low cells. Importantly, there was a strong correlation between [^18^F]FSPG retention and intracellular GSH across all cell lines (R^2^ = 0.89; *p* = 0.0004; **Fig. 1i**), linking [^18^F]FSPG retention to the NRF2-mediated antioxidant program.

### [^18^F]FSPG retention is altered following pharmacological and genetic manipulation of NRF2

To better understand the relationship between NRF2 activity and [^18^F]FSPG retention, we treated NRF2-low cell lines with the KEAP1 inhibitor KI696 (**Fig. 2a**), which interrupts KEAP1/NRF2 binding and stabilizes NRF2^32^. As expected, NRF2 activation by KI696 in NRF2-low cell lines increased the expression (**Fig. 2b**) and activity of system x_c_^−^ (**Fig. 2c**), whilst decreasing intracellular glutamate in H1975 and H23 cells (**Fig. 2d**). Importantly, KI696 treatment significantly raised intracellular GSH concentrations (**Fig. 2e**) and increased [^18^F]FSPG retention in comparison to vehicle controls in all but H1650 cells, where [^18^F]FSPG retention in treated cells was more variable (**Fig. 2f**).

**Figure 2.**
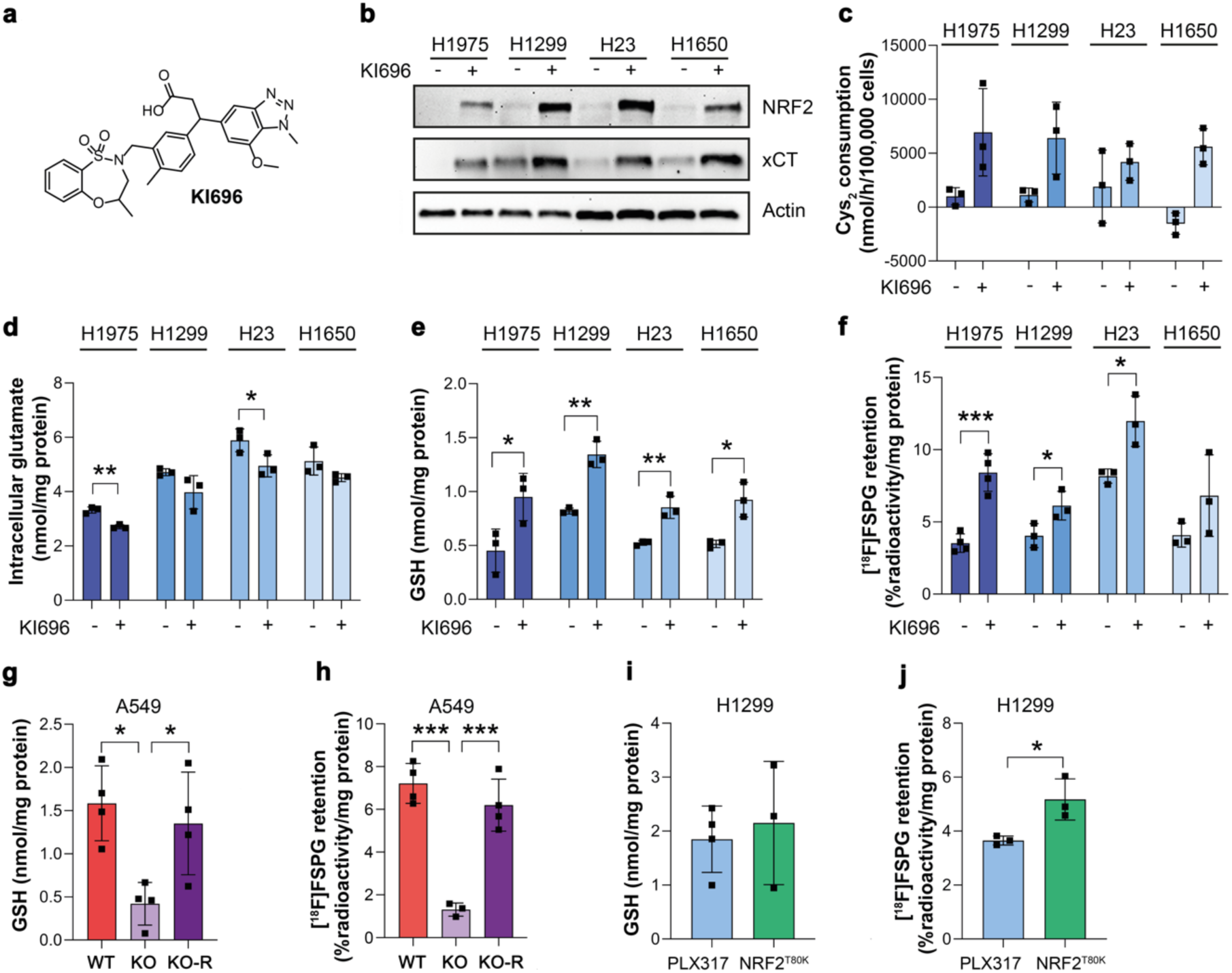
[^18^F]FSPG retention is altered following pharmacological and genetic manipulation of NRF2. a, Chemical structure of KI696. b, Representative western blot of NRF2 and xCT expression in NRF2-low cell lines 24 h post treatment with vehicle control or 200 µM KI696. Actin was used as a loading control. c-f, Analysis of cystine (Cys_2_) consumption (c), intracellular glutamate (d) and intracellular GSH (e) in NRF2-low lines following KI696 treatment compared to vehicle control. f, Intracellular [^18^F]FSPG retention in NRF2-low cells after KI696 treatment compared to vehicle control. g-j. Intracellular GSH (g,i) and [^18^F]FSPG retention (h,j) in genetically modified NSCLC cells. Data are presented as mean ± SD. *, p < 0.05; **; p < 0.01; ***, p < 0.001.

Next, we used A549 (KEAP1 mutant) and H1299 (KEAP1 WT) cells genetically manipulated to alter NRF2 expression levels. We previously reported that knockdown (KO) of NRF2 in A549 cells reduced xCT expression and cystine consumption compared to KO cells restored with ectopic expression of NRF2 (KO-R)^24^. Moreover, introducing the NRF2^T80K^ mutation to H1299 cells increased NRF2 and xCT expression and cystine consumption in H1299 cells compared to empty vector controls (PLX317)^24^. In A549 NRF2 KO cells, GSH was reduced by 73% (*p* = 0.013, *n* = 4) which corresponded to an 82% decrease in [^18^F]FSPG retention compared to WT A549 cells (1.3 ± 0.3% radioactivity/mg protein vs. 7.2 ± 0.9% radioactivity/mg protein, respectively; *n* = 3-4; *p* = 0.0001). Reinsertion of functional NRF2 (A549 NRF2 KO-R) restored GSH levels to baseline (*n* = 4; *p* = 0.75 for A549 vs. A549 KO-R; **Fig. 2g**) and rescued [^18^F]FSPG retention (6.2 ± 1.2% radioactivity/mg protein; *n* = 3; *p* = 0.34; **Fig. 2h**). Conversely, the gain-of-function NRF2 mutation (T80K) in H1299 cells failed to increase intracellular GSH (1.9 ± 0.6 nmol/mg protein in empty vector vs. 2.2 ± 1.1 nmol/mg protein in cells with NRF2^T80K^; *n* = 3-4; *p* = 0.33; **Fig. 2i**). A small but significant increase in [^18^F]FSPG retention, however, was measured in NRF2^T80K^ compared to NRF2^WT^ H1299 cells (5.2 ± 0.8% radioactivity/mg protein vs. 3.7 ± 0.2% radioactivity/mg protein, respectively; *n* = 3; *p* = 0.03; **Fig. 2j**), consistent with our prior finding that NRF2^T80K^ elevates cystine consumption in these cells^24^.

### [^18^F]FSPG PET is a sensitive, non-invasive marker of NRF2 status *in vivo*

To investigate whether [^18^F]FSPG could be used as a non-invasive marker of NRF2 expression *in vivo*, H460 FLuc (NRF2-high) and H1299 FLuc (NRF2-low) NSCLC cell lines were orthotopically grown in the lungs of mice. Tumour progression was longitudinally monitored by bioluminescence imaging (BLI; **Extended Data Fig. 2**) and imaged by [^18^F]FSPG PET when lesions became visible on CT (**Fig. 3a**). Histological analysis of excised lungs revealed multi-focal disease and variable tumour sizes for both H1299 and H460, with substantially higher xCT expression in H460 compared to H1299 tumours (**Fig. 3a**). PET imaging revealed a typical pattern of [^18^F]FSPG distribution, characterised by low physiological uptake in all healthy organs except the pancreas, and elimination via the urinary tract (**Fig. 3b** and **Extended Data Fig. 3**). [^18^F]FSPG retention in both tumours was clearly visible above background (**Fig. 3b**), with image-quantified [^18^F]FSPG retention ∼3-fold higher in H460 tumours compared to H1299 tumours (8.3 ± 1.6% ID/g protein vs. 2.8 ± 1.5% ID/g protein; *n* = 7-21 lesions from 3-9 mice; *p* < 0.0001; **Fig. 3c**). NRF2 was substantially higher in H460 lesions compared to H1299; a pattern that was replicated with xCT (**Fig. 3d**).

**Figure 3.**
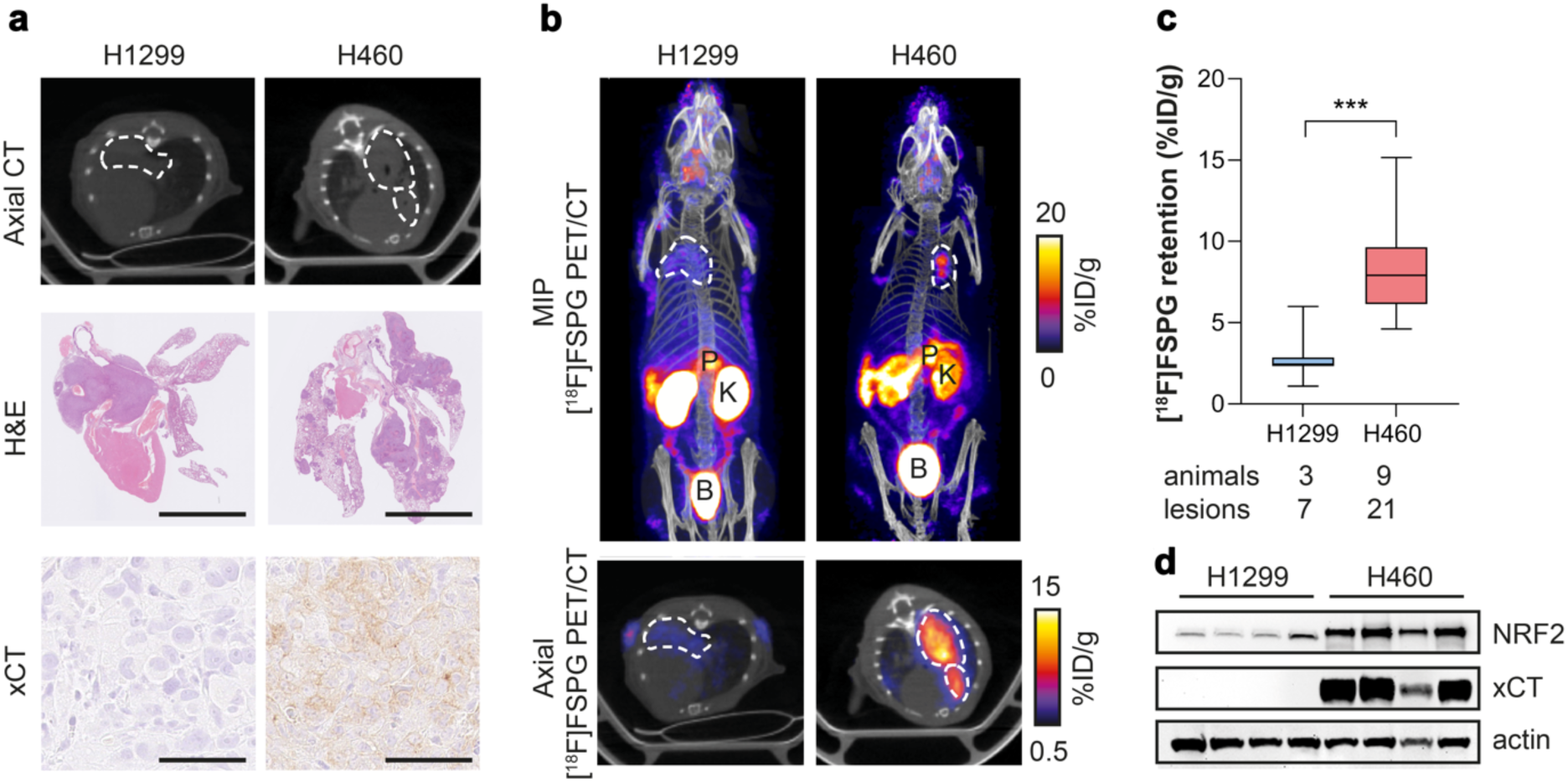
[^18^F]FSPG PET can differentiate NRF2-high from NRF2-low tumours when grown orthotopically in the lungs of mice. a, Single slice CT axial images (top) and ex vivo H&E images (middle; scale bar = 5 mm) of lungs containing H1299 or H460 tumours, with xCT staining of corresponding tumours (bottom; scale bar = 50 µm). b, Representative in vivo [^18^F]FSPG PET/CT maximum intensity projections (MIPs; top) and axial single-slice PET/CT (bottom) of mice bearing H1299 or H460 orthotopic lung tumours. Dashed lines represent the tumour outline. c, Quantified [^18^F]FSPG retention in individual tumour lesions from orthotopic tumour-bearing mice. d, Representative western blot for xCT and NRF2 expression in H1299 and H460 orthotopically grown tumours (n = 4 per tumour). Actin was used as a loading control.

### [^18^F]FSPG retention is increased in tumours of genetically engineered mice with Nrf2 activation

To examine whether [^18^F]FSPG PET could identify enhanced Nrf2 activity in lung tumours of immunocompetent mice, a conditional knock-in mouse model of Nrf2^D29H/+^ was used (**Fig 4a**). Lung tumours were induced following intra-nasal administration of adenoviral-Cre in the Kras^G12D/+^; p53^flox/flox^ (KP) and Kras^G12D/+^; p53^flox/flox^; Nrf2^D29H/+^ (KPN) model^28^. Viral infection of the lungs of KP and KPN mice resulted in the development of tumours after approximately three months, with multiple lesions visible by CT (**Fig. 4b**). Interestingly, although the number of tumours arising from KPN mice were greater than those of KP mice (105 vs. 63), the total tumour volumes between cohorts were similar, at 22.2 ± 9.4% and 18.1 ± 6.5% of total lung volume, respectively (*n* = 5-6; *p* = 0.44; **Fig. 4c**). We next performed PET imaging with KP and KPN mice to non-invasively profile tumour-associated Nrf2 activity with [^18^F]FSPG. Representative single slice coronal [^18^F]FSPG PET/CT images of 40-60 min summed radioactivity are shown in **Fig. 4d**. [^18^F]FSPG retention was high in KP tumours (10.1 ± 3.3% ID/g; *n* = 62 lesions), which was further increased on average 3.6-fold in KPN tumours (36.8 ± 14.3% ID/g; *n* = 104 lesions; *p* < 0.0001; **Fig. 4e**). There was a heterogenous distribution of [^18^F]FSPG retention across KPN lesions, with some tumours reaching >80% ID/g. Following imaging, tumours harvested from KP and KPN mice confirmed elevated Nrf2 in tumours expressing heterozygous Nrf2^D29H/+^ (**Fig. 4f**). As expected, the Nrf2^D29H/+^ mutation also increased xCT compared to Nrf2^WT^ tumours (**Fig. 4g,h**). We previously reported that Nrf2 and the Nrf2-regulated gene Nqo1 are downregulated in high-grade tumours that arise from Keap1/Nrf2 mutant models^28^. Similarly, xCT expression was reduced in the KPN model as tumours progressed from hyperplasia (atypical adenomatous and bronchiolar) and low-grade tumours to higher grades (**Fig. 4i**).

**Figure 4.**
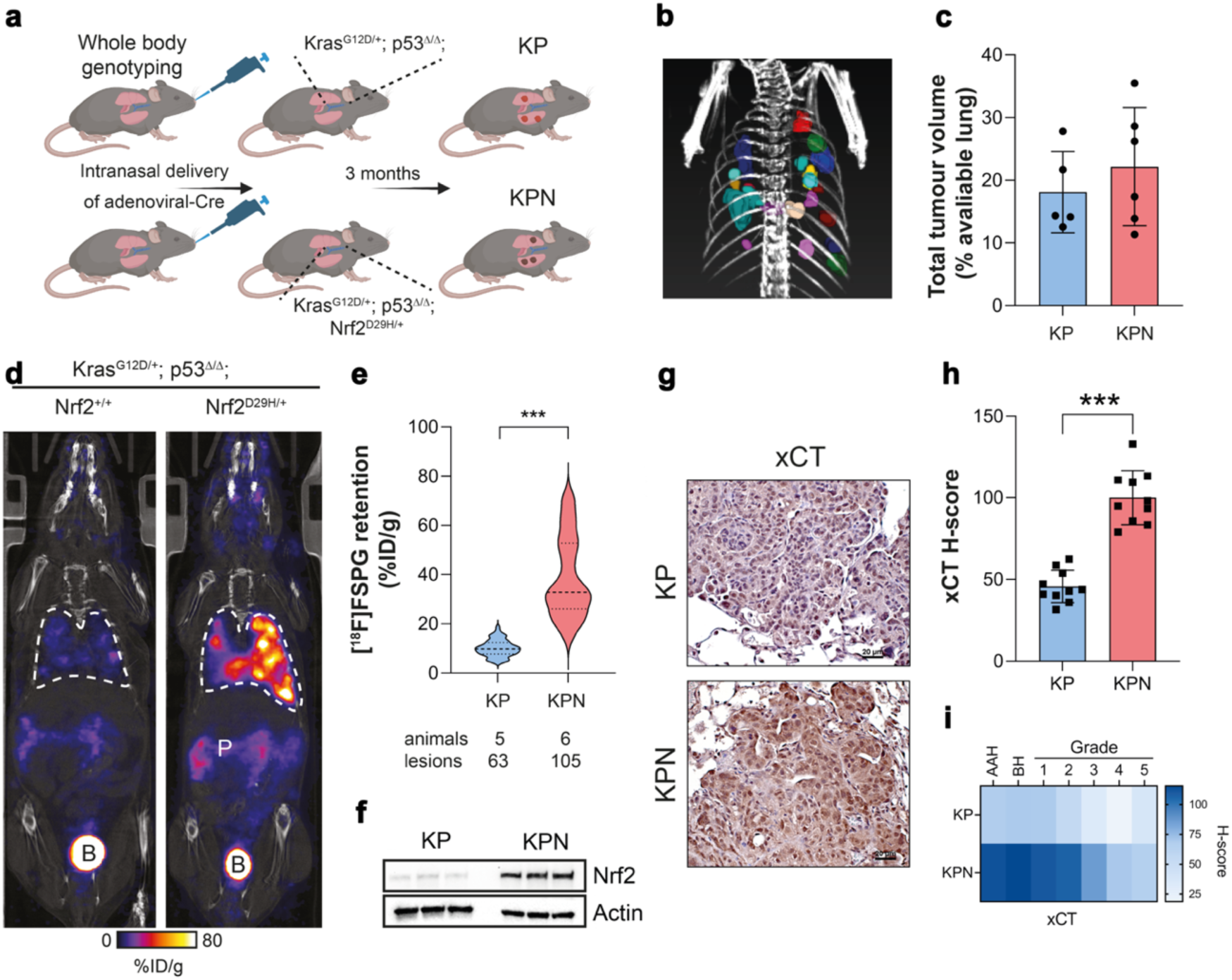
[^18^F]FSPG retention is increased in Nrf2 mutant mice. a, Scheme depicting tumour formation in KP and KPN mice. KP mice conditionally express oncogenic Kras and have loss of p53 function. KPN mice conditionally express oncogenic Kras, have loss of p53 function and express a mutant Nrf2 which increases Nrf2 protein levels. b, CT MIP representing individual 3D tumour regions of interest. c, Total tumour volumes in the lungs of KP and KPN mice. Scatter plots represent individual animals. d, Representative coronal [^18^F]FSPG PET/CT images of 40-60 min summed activity in KP and KPN tumour-bearing mice. Dashed white lines indicate the lung. B, bladder; P, pancreas. e, Violin plots of [^18^F]FSPG tumour retention from individual lesions. n = 63-105 lesions from 5-6 mice per cohort. f, Nrf2 expression in KP and KPN tumour lesions. Actin was used as a loading control. g, Representative IHC staining of xCT from lesions taken from KP and KPN mice (scale bars = 20 μM). h, H-scores for xCT IHC staining. i, Heatmap depicting the H-scores for xCT IHC staining by tumour grade. AAH, adenomatous atypical hyperplasia; BH, bronchiolar hyperplasia. ***, p < 0.001.

### Elevated [^18^F]FSPG retention in NRF2-mutant tumours is recapitulated in patient-derived xenografts

Lung TRACERx (TRAcking Cancer Evolution through therapy (Rx)) is a prospective cohort study that aims to define how intratumour heterogeneity affects the risk of reoccurrence and survival in NSCLC^33^. Transcriptomic data from the TRACERx 421 cohort^29^ revealed highly variable *NFE2L2* and NRF2-regulated gene mRNA abundance, both within and between tumours (**Extended Data Fig. 4**). There was good concordance between *SLC7A11* (which encodes xCT), *NQO1*, *GCLC* and *GCLM* median expression levels across tumours (**Fig. 5a**), providing an ‘antioxidant signature’ associated with the NRF2 transcriptional program. We identified patient-derived xenograft (PDX) models from TRACERx patients^30^ which were either *NFE2L2* wildtype (WT; CRUK0640 region 8; R8) or contained truncal alterations in *NFE2L2* (c.G187C/p.E63Q and c.G352C/p.E118Q; CRUK0772 R1) ^30^. CRUK0640 R8 had truncal mutations in *NF1* and *KMT2D*, while CRUK0772 R1 had *TP53* and *RB1* mutations in addition to the *NFE2L2* mutations. In patient RNA sequencing data, *SLC7A11*, *NQO1*, *NFE2L2* and *GCLM* gene expression was higher in CRUK0772 tumour regions compared to CRUK0640 (**Extended Data Fig. 5**).

**Figure 5.**
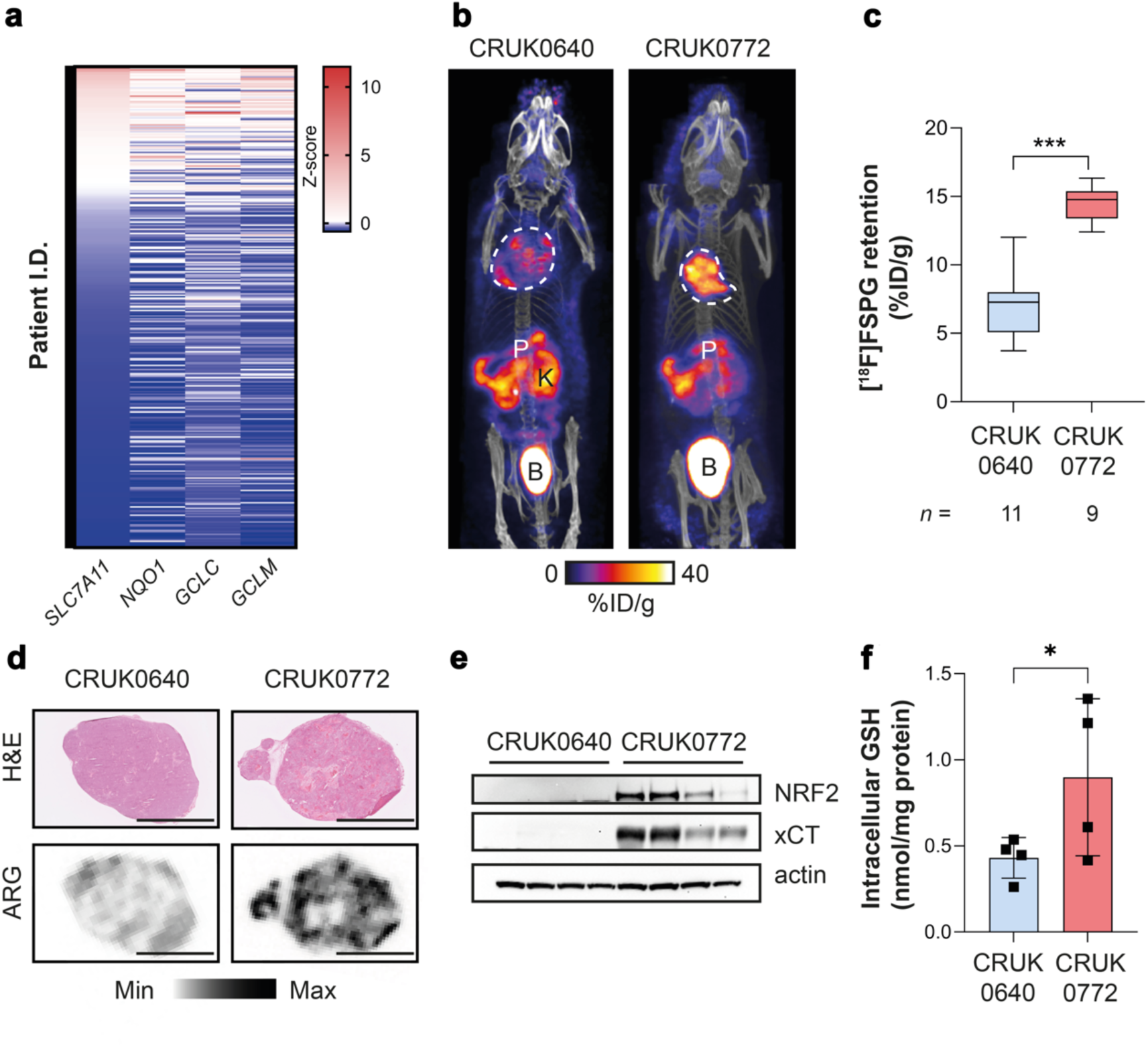
An antioxidant gene signature accompanies NRF2 mutations in patient tumours and patient derived xenograft (PDX) models, which is detectable by [^18^F]FSPG PET. a, Expression of NRF2-regulated genes in the TRACERx 421 patient cohort. b, Representative [^18^F]FSPG MIP of mice bearing PDXs either with (CRUK0772 R1) or without (CRUK0640 R8) a NRF2 mutation. c, Quantification of [^18^F]FSPG tumour retention (n = 9-11 mice/group). d, Post-imaging autoradiograms (ARG) from PDXs, illustrating the intratumoural heterogeneity of [^18^F]FSPG retention. Scale bar = 5 mm. e, xCT and NRF2 protein expression in PDX xenografts (n = 4 per tumour type). Actin was used as a loading control. f, GSH measurements from PDX tumours (n = 4). *, p < 0.05; ***, p < 0.001.

Once tumour xenografts reached 150 mm^3^, mice bearing NRF2 WT and mutant tumours underwent [^18^F]FSPG PET/CT imaging. As with orthotopic and syngeneic tumours, [^18^F]FSPG retention was higher in the NRF2-mutant xenografts compared to WT (**Fig. 5b** and **Extended Data Fig. 6**), at 14.8 ± 2.8% ID/g and 7.0 ± 2.4% ID/g, respectively (*n* = 9-11; *p* < 0.0001; **Fig. 5c**). There was a high degree of intratumoural heterogeneity of [^18^F]FSPG retention for both PDX models which was not related to cellularity, as confirmed by *ex vivo* autoradiography (**Fig. 5d**). Regional sampling of the xenograft tissue revealed high but variable xCT expression in CRUK0772 R1 xenografts that matched the pattern of NRF2 expression, whereas neither xCT nor NRF2 could be detected by western blot in CRUK0640 R8 xenografts (**Fig. 5e**). The elevated antioxidant capacity of NRF2-mutant xenografts was reflected in a doubling of tumoural GSH (0.90 ± 0.46 nmol/mg protein for CRUK0772 R1 compared to 0.43 ± 0.12 nmol/mg protein for CRUK0640 R8; *n* = 4; *p* = 0.047; **Fig. 5f**), again recapitulating the pattern of [^18^F]FSPG retention.

### System x_c_^−^ is a vulnerability that can be exploited for targeted therapy

Antibody-drug conjugates (ADCs) are a promising class of cancer therapeutics that provides precision targeting of cell surface antigens and delivery of a therapeutic payload following receptor internalisation. Given that xCT is upregulated in tumours with activated NRF2, xCT may provide a specific vulnerability of therapy-resistant cancer that can be exploited therapeutically. To test this hypothesis, we assessed the efficacy of a humanised xCT-targeting monoclonal antibody conjugated to tesirine (HM30-tesirine; **Fig. 6a**) in NSCLC. Conjugation of tesirine to the antibody had no effect on its stability (**Extended Data Fig. 7**) and HM30-tesirine selectively bound to xCT present in the lysates of H460 cells, whereas no binding was present in low xCT-expressing H1299 cells (**Fig. 6b**). HM30-tesirine induced a dose-dependent increase in cell kill in H460 cells, with an EC_50_ of 3.7 nM ± 2.4 nM, whereas the NRF2 and xCT-low H1299 cells had an EC_50_ of 120 nM ± 50 nM, suggesting reduced sensitivity to this ADC (*n* = 3; *p* = 0.018; **Fig. 6c**).

**Figure 6.**
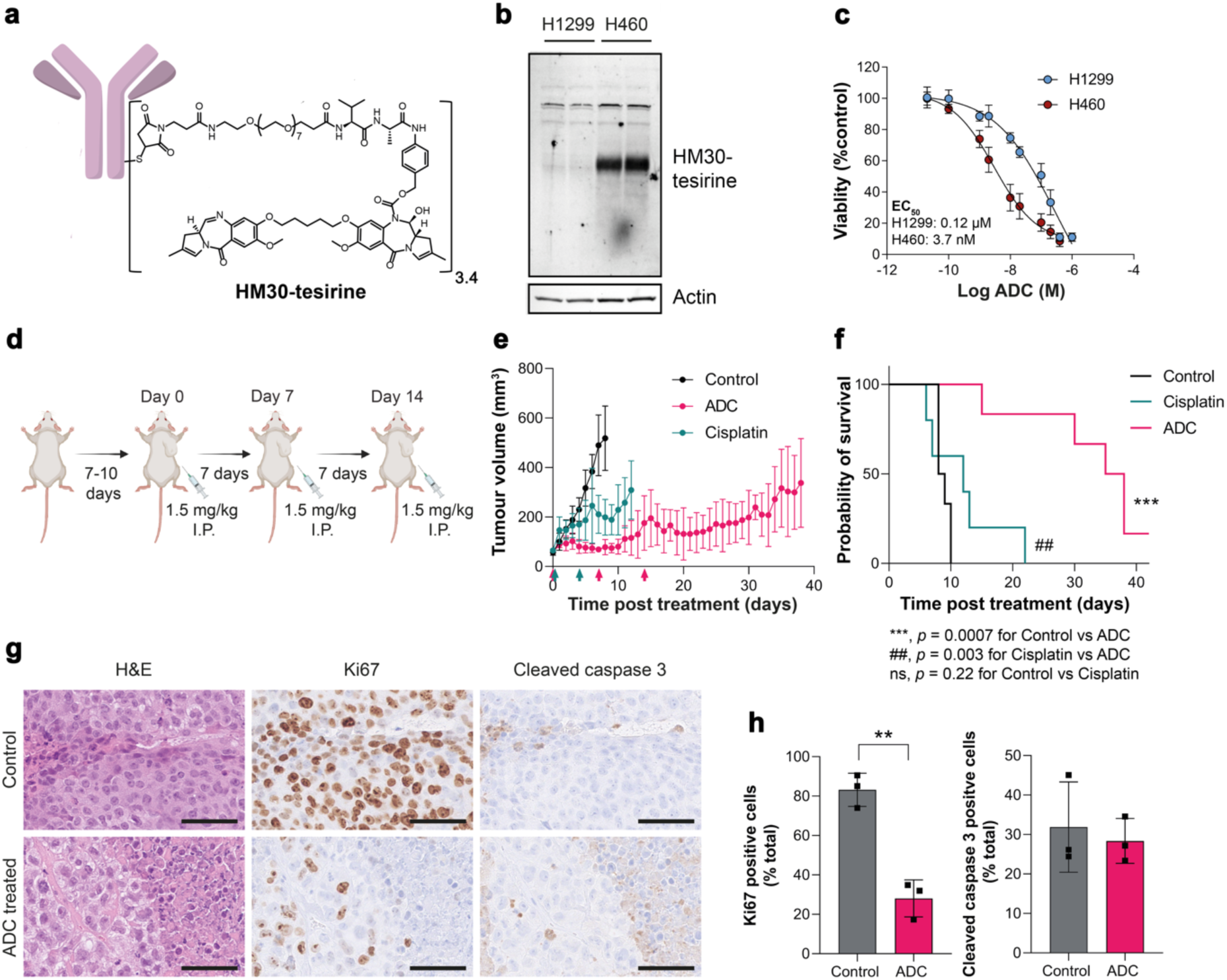
An xCT-ADC controls tumour growth and prolongs survival of mice bearing H460 tumours. a, Structure of the anti-xCT tesirine conjugate, HM30-tesirine. b, Western blot using HM30-tesirine as the primary antibody in H1299 and H460 cell lysates. Actin was used as a loading control. c, HM30-tesirine MTT dose-response in H460 and H1299 cells. d, Schematic representation of HM30-tesirine treatment course in balb/c nu/nu mice. Antitumour activity (e) and survival benefit (f) of control (saline treated), cisplatin treated, and HM30-tesirine treated mice. For statistical analysis, a two-tailed t-test (for tumour volume) and a log-rank test (for survival curve) were used. To control the family-wise error rate in multiple comparisons, crude p-values were adjusted by the Holm–Bonferroni method. h, IHC for Ki67 and cleaved caspase 3 from FFPE tumours taken at endpoint. Scale bar, 50 µm. h, Corresponding quantification of tissue staining. Data are presented as the mean values ± SD (n = 3).

Next, the *in vivo* efficacy of HM30-tesirine was examined in mice bearing subcutaneous H460 tumours. Tumours were grown until they reached ∼70 mm^3^, approximately 7-10 days post inoculation, before being randomised into vehicle, cisplatin, and HM30-tesirine treatment groups, with mice receiving three intraperitoneal doses of 1.5 mg/kg HM30-tesirine seven days apart (**Fig. 6d**). There was no acute toxicity associated with HM30-tesirine administration, assessed through measurements of body weight (**Extended Data Fig. 8**) and observation of clinical symptoms. HM30-tesirine substantially inhibited tumour growth compared to vehicle and cisplatin-treated mice over the six-week treatment study, with effects still present >20 days after the last course of treatment (**Fig. 6e**). The initial dose of HM30-tesirine maintained tumour volumes close to their pre-treatment size eight days post-treatment (78 ± 31 mm^3^), whereas control and cisplatin-treated tumours had reached 518 ± 130 mm^3^ and 199 ± 72 mm^3^, respectively by this time (*n* = 3-6; *p* = 0.0013 for control vs. cisplatin; *p* <0.0001 for control vs. HM30-tesirine; *p* = 0.1 for cisplatin vs. HM30-tesirine). HM30-tesirine treatment extended the lives of all mice enrolled, increasing the median survival from 8.5 days in control mice to 36.5 days following HM30-tesirine treatment (*p* = 0.0007; **Fig. 6f**). Conversely, whilst cisplatin induced a small but significant reduction in rate of tumour growth, this standard-of-care treatment had no effect on overall survival compared to control tumours (median survival = 12 days; *p* = 0.22 for control vs. cisplatin).

Immunohistochemical analysis of tumours taken at humane endpoints or at the end of the six-week study revealed a 65% reduction in tumour cell proliferation in mice treated with HM30-tesirine, with 28 ± 9.4% of cells staining positive for Ki67, compared to 83 ± 8.4% in control tumours (*n* = 3 individual tumours; *p* = 0.0016; **Fig. 6g,h**). At these late time points, there was no difference in levels of apoptosis between groups, as shown by cleaved caspase 3 staining (32 ± 11% positive cells for control vs. 28 ± 5.7% for HM30-tesirine; *n* = 3; *p* = 0.66; **Fig. 6g,h**).

## Discussion

Resistance to therapy is one of the biggest problems in clinical oncology. Despite a revolution in new anti-cancer therapies, such as check-point inhibitors and proton beam therapy, durable responses are often not observed due to acquired or innate resistance to existing treatment regimens^34, 35^. Currently, there is no satisfactory way to identify patients that are refractive to therapy early on in their treatment pathway^36^. Treatment failure plays a critical role in the management of patients with NSCLC, where survival rates have struggled to improve over the last ten years despite advances in prevention, screening and treatment^37^. In NSCLC, constitutive NRF2 activation results in resistance across the spectrum of currently available therapeutics^9–12^. A non-invasive measure of NRF2 activation may therefore provide an attractive solution for the prediction of therapy resistance in NSCLC, which may further reveal cancer-specific vulnerabilities for the precision treatment of refractive disease.

Using a combination of metabolomics, genetic engineering, drug treatment, and biochemical analyses, we comprehensively demonstrated the ability of system x_c_^−^ to report on NRF2 activation. In NSCLC cells grown in culture, KEAP1 loss-of-function mutations significantly increased xCT expression, had higher cystine consumption from the medium, and decreased intracellular glutamate concentrations – features indicative of increased cystine/glutamate exchange by system x_c_^−^. Given that cystine influx drives GSH biosynthesis, it was unsurprising that NRF2-high cells had elevated steady-state GSH and a concomitant reduction in ROS. Similar observations were made when KEAP1 WT cells were treated with the KEAP1/NRF2 inhibitor KI696: xCT was increased, along with GSH concentration and cystine utilisation. NRF2 depletion in KEAP1 mutant A549 cells has the opposite effect, which was rescued following ectopic expression of NRF2.

[^18^F]FSPG PET has been used to image system x_c_^−^ activity in a range of human malignancies, including NSCLC^38–41^. Although exclusively used in clinical trials as a diagnostic agent, we^21–23^ and others^42^ have shown that [^18^F]FSPG is a sensitive marker of therapy-induced changes in tumour redox status. Here, disruption of redox homeostasis through NRF2 activation increased [^18^F]FSPG cell retention; changes that were tightly correlated to cellular GSH concentration (R^2^ = 0.89). Interestingly, whilst the R272L NRF2 mutation in H1944 cells increased both NRF2 and xCT expression, increased GSH and [^18^F]FSPG retention compared to WT cells was muted, suggesting that not just system x_c_^−^ activity, but the fate of its substrates is reflected in the imaging readouts. In our genetic models and following pharmacological intervention, [^18^F]FSPG retention mirrored NRF2 activation levels.

Whilst other non-radioactive assays are better suited to evaluate NRF2 status in isolated cells, PET imaging facilitates the measurement of biochemical processes across the whole body, permitting assessment of the entire tumour burden^43^. This is of particular importance when evaluating clonal heterogeneity, as was evident in our PDX model studies (**Fig. 5**). The TRACERx consortium has pioneered multi-region tumour sampling and phenotyping using next-generation sequencing approaches, revealing new insights into tumour evolution^33^. However, this extensive tumour characterisation, using specialised clinical and data analysis pipelines, is currently limited to a minority of patients in research studies. Alternatively, [^18^F]FSPG imaging could be made available to patients at most major hospitals which currently use [^18^F]2-fluoro-2-deoxy-D-glucose ([^18^F]FDG) for tumour staging/restaging. Across a range of genetically engineered, patient-derived, and orthoptic NSCLC models, [^18^F]FSPG retention was increased in tumours with NRF2 activation compared to tumours with a normal-functioning NRF2/KEAP1 axis. The heterogeneity of this imaging signal was reflected in *ex vivo* markers of NRF2, xCT, and GSH, providing further evidence that [^18^F]FSPG reports on tumour redox status. Importantly, RNA sequencing data from TRACERx patients suggested that a ‘redox signature’ of NRF2-regulated genes that could be used to select patients for further monitoring by [^18^F]FSPG PET imaging.

Given that NRF2 and xCT are intrinsically interconnected, and NRF2 confers therapy resistance, we investigated whether xCT was a vulnerability that could be exploited for the treatment of refractive disease. We assessed an xCT-specific ADC, HM30-tesirine, in mice bearing H460 tumours which harbour the D236H KEAP1 loss-of-function mutation. H460 tumours are intrinsically therapy-resistant, due in-part to NRF2 activation^44^, with cisplatin treatment yielding only a moderate reduction in tumour growth rate. Conversely, HM30-tesirine treatment resulted in sustained tumour growth suppression and decreased proliferation, which was maintained following removal of treatment, likely due to the long circulation half-life of the ADC. Importantly, HM30-tesirine significantly extended the lives of H460 tumour-bearing mice compared to both cisplatin- and vehicle-treated animals. Small molecule drugs that target xCT, such as imidazole ketone erastin and sorafenib, have also displayed potent anti-tumour effects^45^. However, the relatively poor metabolic stability of these therapeutics^46^, and additional mechanisms of action besides xCT inhibition^47^, may preclude their future clinical utility. This first-in-class xCT ADC may circumvent some of these issues, providing a new therapeutic avenue for the treatment of refractive NSCLC.

Despite these advances, the imaging and targeted treatment of NSCLC with high system x_c_^−^ expression holds specific challenges. [^18^F]FSPG is bidirectionally transported across the plasma membrane. As shown here, tumours with high NRF2 not only have elevated uptake, but also increased efflux from the cell (**Fig. 1h**, **Extended Data** Fig. 1). Consequently, net [^18^F]FSPG in NRF2 activated tumours was increased ∼3-fold compared to NRF2 WT tumours, despite orders of magnitude differences in protein expression. Moreover, the activity of system x_c_^−^ is not solely driven by transporter expression, but by the concentrations of both cystine and glutamate across the membrane^48^. Nutrient composition in the tumour microenvironment, or metabolic alterations (e.g. anaplerosis), may therefore affect [^18^F]FSPG irrespective of NRF2 status. Antibody-based approaches (such as HM30-tesirine) are not dependent on transporter activity and may therefore simplify therapeutic strategies. System x_c_^−^ however is not exclusively expressed on tumour cells, with high expression found in the brain, pancreas, and components of the immune system^49, 50^. Although HM30-tesirine was well tolerated in mice, patients receiving xCT-targeted therapies should be monitored for on-target, off-tumour toxicity. As with any other surrogate marker, system x_c_^−^ expression is also influenced by factors other than NRF2, such as ATF4, p53, and CD44v^51^. Consequently, prospective clinical trials are required to assess the specificity of [^18^F]FSPG for NRF2 in humans and to determine the thresholds of [^18^F]FSPG retention to classify tumours into NRF2-high or NRF2-low.

In summary, we describe the use of a clinically-tested PET radiotracer, [^18^F]FSPG, for the imaging of NRF2 activation in NSCLC. This study sets the foundation for the clinical assessment of NRF2 in NSCLC patients with [^18^F]FSPG at King’s College London (clinical trials number: NCT05889312). If successful, the combined imaging and xCT-targeted treatment of NSCLC with activating NRF2/KEAP1 mutations may represent a new paradigm for patients with therapy-resistant disease.

## Acknowledgements

The authors thank members of the lung TRACERx consortium, in particular Mariam Jamal-Hanjani and Crispin Hiley. We are grateful for the support provided by Tammy Kalber and Stephen Patrick who facilitated the GEMM imaging studies at UCL, and both Kavitha Sunassee and Jana Kim, who supported the imaging studies at KCL. Finally, we thank Les Steward and Joseph Patti from AgilVax for insightful discussions and for providing HM30-tesirine.

