## Extended Figures for "Imaging the master regulator of the antioxidant response in non-small cell lung cancer with positron emission tomography"

**EXTENDED FIGURES AND LEGENDS**

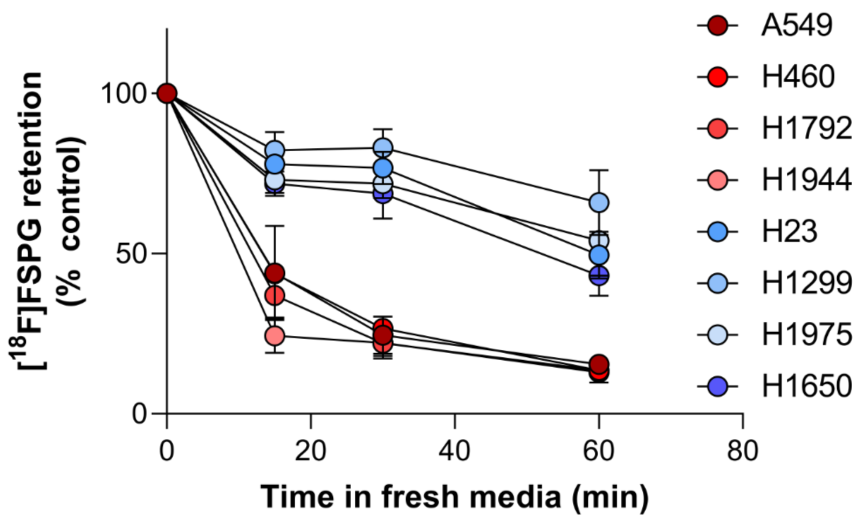


**Extended Figure 1.** Time course of [$^{18}$F]FSPG efflux across NRF2-high (red) and NRF2-low NSCLC cells (blue).

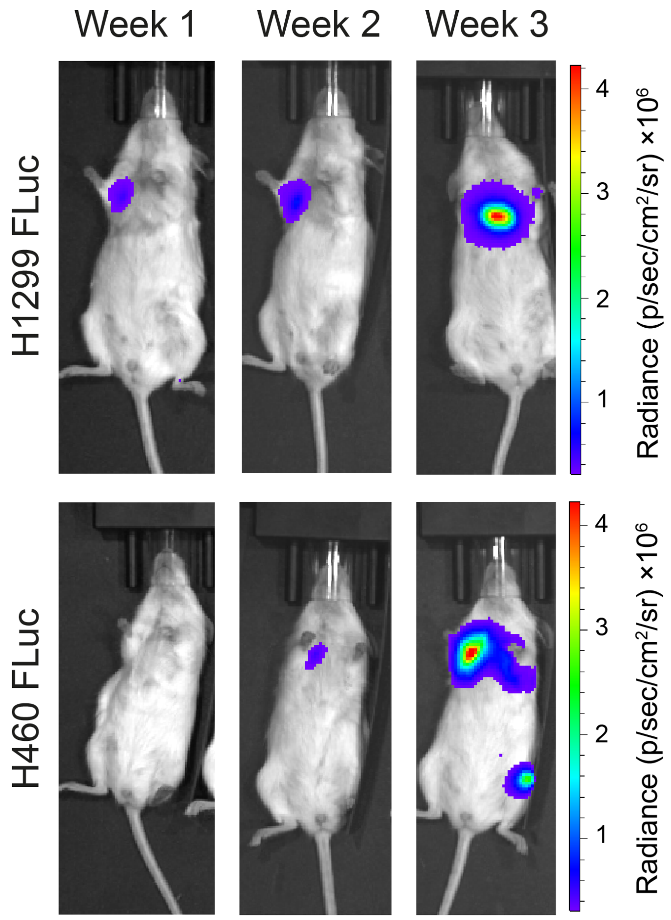


**Extended Figure 2.** Representative BLI from mice orthotopically implanted with H460 FLuc and H1299 FLuc tumour cells.

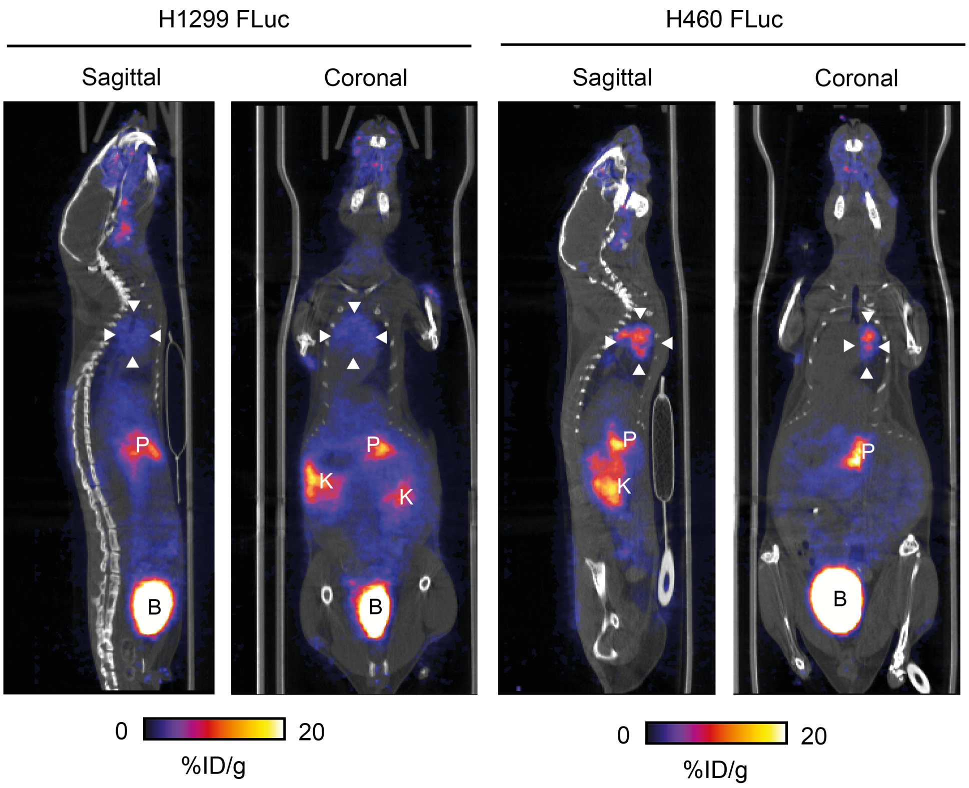


**Extended Figure 3.** Representative single slice PET/CT images from mice orthotopically implanted with H460 FLuc and H1299 FLuc tumour cells. Arrowheads denote the tumour. K, kidney; P, pancreas; B, bladder.

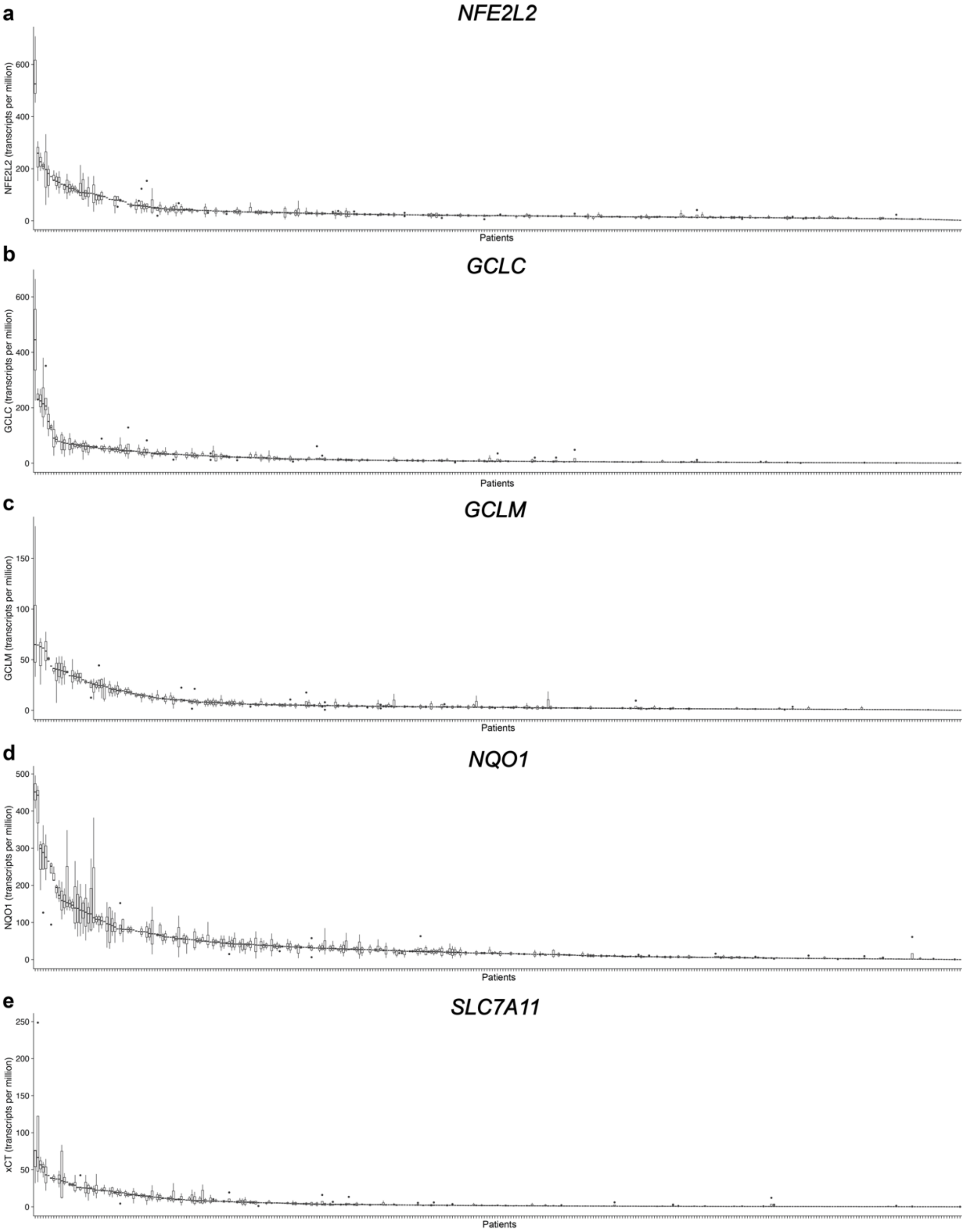


**Extended Figure 4.** Multi-region bulk RNA sequencing data from 882 samples from 344 NSCLC patients. Plots show expression (transcripts per million) of *NFE2L2* (a), *GCLC* (b), *GCLM* (c), *NQO1* (d) and *SLC7A11* (e). Box plots show the lower and upper quartiles (box limits) and the median (centre line). The length of whiskers is 1.5 × Interquartile range and outlier points outside this range are shown. Patients are ordered from highest to lowest median expression of each gene along the x-axis, and thus are not in the same order between plots.

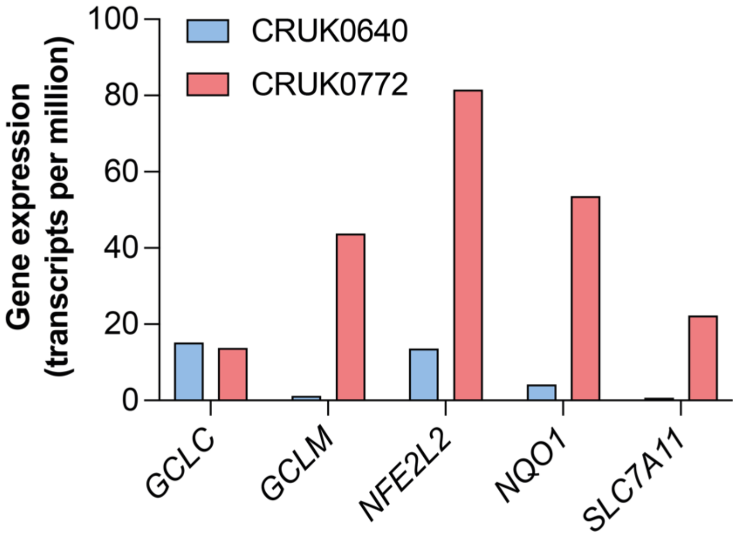


**Extended Figure 5.** Mean NRF2-regulated gene expression in CRUK0640 and CRUK0772 tumour resection samples from TRACERx421 RNA sequencing data.

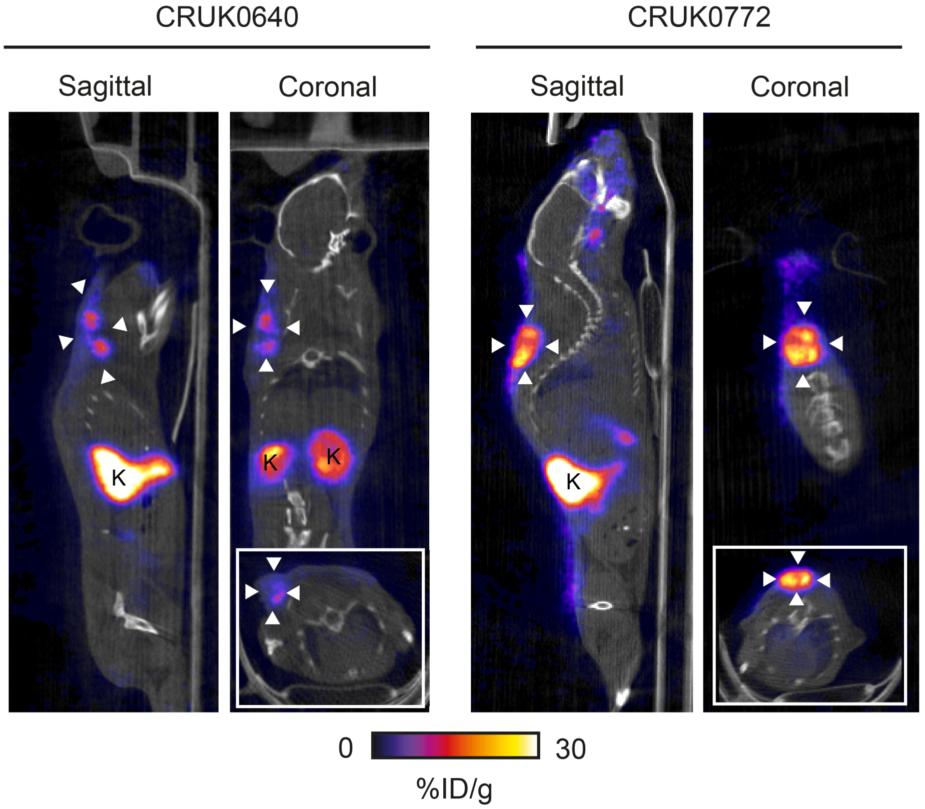


**Extended Figure 6.** Representative single slice PET/CT images from mice bearing CRUK0640 or CRUK0772 patient-derived xenografts. Arrowheads denote the tumour. K, kidney.

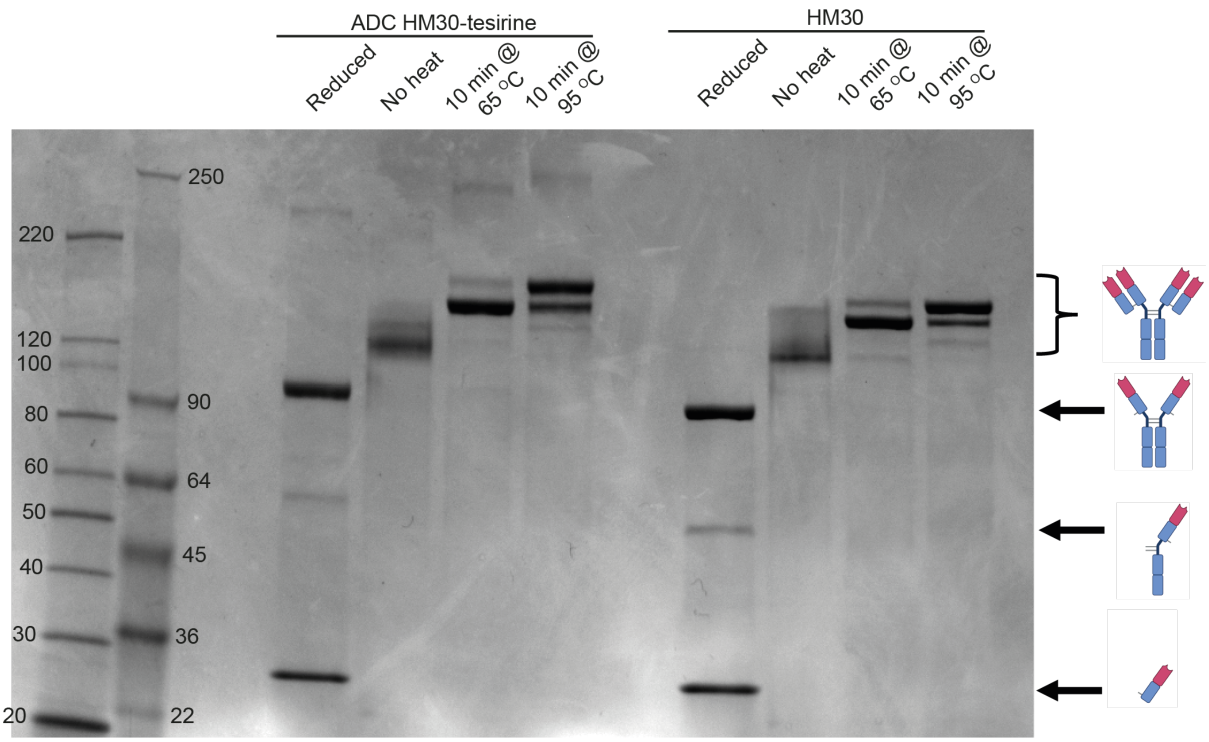


**Extended Figure 7.** HM30-tesirine and unconjugated HM30 monoclonal antibody SDS-PAGE profile with a range of pre-treatment conditions.

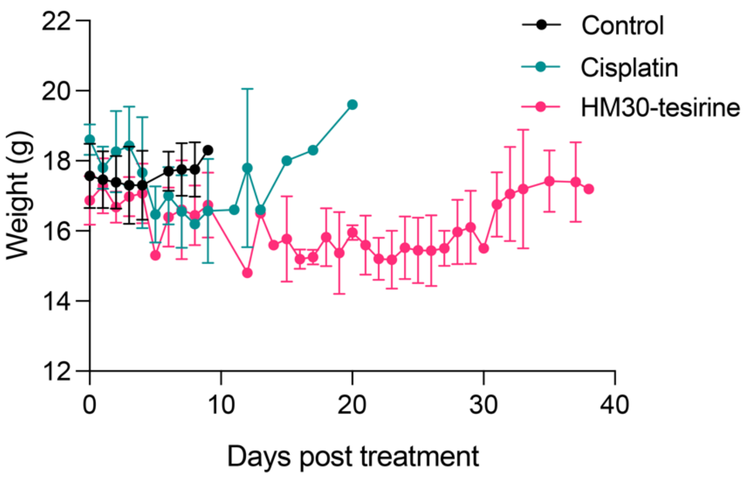


**Extended Figure 8.** Body weight from H460 tumour-bearing mice following treatment with vehicle control, cisplatin, and HM30-tesirine.
